# Chromosomally unstable tumor cells specifically require KIF18A for proliferation

**DOI:** 10.1101/2020.06.18.159327

**Authors:** Carolyn Marquis, Cindy L. Fonseca, Katelyn A. Queen, Lisa Wood, Sarah E. Vandal, Heidi L.H. Malaby, Joseph E. Clayton, Jason Stumpff

## Abstract

Chromosomal instability (CIN), characterized by frequent missegregation of chromosomes during mitosis, is a hallmark of tumor cells caused by changes in the dynamics and control of microtubules that comprise the mitotic spindle^1–3^. Thus, CIN tumor cells may respond differently than normal diploid cells to treatments that target mitotic spindle regulation. We tested this idea by inhibiting a subset of kinesin motor proteins that control spindle microtubule dynamics and mechanics but are not required for the proliferation of near-diploid cells. Our results indicated that KIF18A was required for proliferation of CIN cells derived from triple negative breast cancer or colorectal cancer tumors but was not required in near-diploid cells. CIN tumor cells exhibited mitotic delays, multipolar spindles due to centrosome fragmentation, and increased cell death following inhibition of KIF18A. Sensitivity to KIF18A knockdown was strongly correlated with centrosome fragmentation, which required dynamic microtubules but did not depend on bipolar spindle formation or mitotic arrest. Our results indicate the altered spindle microtubule dynamics characteristic of CIN tumor cells can be exploited to reduce the proliferative capacity of CIN cells.

## Introduction

Genetic instability is a common feature of tumor cells, and a large number of tumor cells exhibit frequent loss or gain of chromosomes^1^. This chromosomal instability (CIN) is primarily attributable to defects leading to abnormal interactions between chromosomes and mitotic spindle microtubules, which in turn increase chromosome segregation errors^2–7^. While CIN contributes to tumor progression, heterogeneity, drug resistance, and metastasis, it has been proposed that the same properties driving instability could provide an Achilles’ heel for CIN cell-specific targeted therapies^1,8,9^. Compared to chromosomally stable cells, CIN cells display increased spindle microtubule polymerization and reduced turnover of the attachments between spindle microtubules and kinetochores, which are specialized protein structures that assemble at the centromeric regions of mitotic chromosomes^2,3^. Thus, CIN cells may be particularly vulnerable to anti-mitotic therapies that target the microtubule cytoskeleton.

Consistent with this idea, microtubule-targeting agents are effective therapeutics for a wide variety of tumors^10^. Paclitaxel, a microtubule stabilizing drug routinely utilized to treat solid tumors, was originally demonstrated to induce cytotoxicity by preventing cells from completing mitosis^11^. However, due to adverse side effects associated with the broad inhibition of microtubule function, significant effort has been made to identify mitotic regulators that could be targeted with lower toxicity in cancer patients. While drugs targeting proteins essential for mitosis have shown promise in preclinical models, they have been largely unsuccessful in clinical trials^12^. One explanation for the apparent paradox presented by failed mitotic targeting strategies and the effective therapeutic results seen with paclitaxel is that paclitaxel may not kill tumor cells in vivo simply by preventing mitotic progression. This idea is supported by work demonstrating that clinically relevant doses of paclitaxel induce abnormal, multipolar divisions in tumors, rather than preventing mitotic division altogether^11,13^. Furthermore, paclitaxel leads to induction of micronuclei due to chromosome segregation errors, which may activate innate immune pathways^14^. Thus, efforts to mimic the effects of paclitaxel on mitotic cells need to be refocused towards identifying proteins that can be targeted to disrupt normal bipolar divisions, ideally in a tumor cell specific manner.

Here, we tested the hypothesis that altered mitotic microtubule dynamics in CIN cells may confer sensitivity to inhibition of proteins that regulate microtubule dynamics or generate forces within mitotic spindles. Ideal targets would reduce CIN cell proliferation by inducing mitotic defects specifically in tumor cells. We focused our efforts on kinesin motors known to regulate spindle microtubule dynamics and mechanics that are also largely dispensable for division in diploid somatic cells.

## Results

### KIF18A is required for the proliferation of CIN tumor cells

To compare the impacts of altered kinesin function in cells with or without CIN, we measured cell proliferation in both stable, diploid breast epithelial MCF10A cells and the chromosomally unstable triple negative breast cancer (TNBC) cell lines MDA-MB-231, MDA-MB-468, and HCC1806^15^ following knockdown (KD) of kinesin motor proteins. Specifically, the effects of KIF18A, KIF18B, KIF4A, KIF22/KID, and KIF2C/MCAK KD were determined (Extended Data Fig 1). Cell proliferation was measured using an automated, high-contrast brightfield microscopy-based kinetic assay (Extended Data Fig 2). KIF18A KD significantly reduced proliferation of all three TNBC cell lines, but did not affect the growth of diploid MCF10A cells (Fig 1 A-B). To determine if this trend holds in other tumor cell types, we measured proliferation in colorectal cancer (CRC) cells categorized as displaying either chromosomal instability (CIN) or microsatellite instability (MSI), a form of genomic instability arising from defective DNA repair in near-diploid tumor cells^16^. KIF18A KD significantly reduced the proliferation of two CIN cell lines but had minor effects on the proliferation of MSI cells (Fig 1C, Extended Data Fig 1). The proliferation of chromosomally unstable HeLa Kyoto cells, which have been used extensively for studies of KIF18A function, was also reduced by KIF18A KD (Fig 1C). CIN cells sensitive to KIF18A knockdown exhibited increased cell death following KIF18A siRNA treatment, while near-diploid HCT116 and MCF10A cells did not (Extended Data Fig 3). Taken together, the diploid cells tested here did not require KIF18A to proliferate, while the majority of CIN tumor cells displayed a dependence on KIF18A for efficient growth and survival.

**Figure 1.**
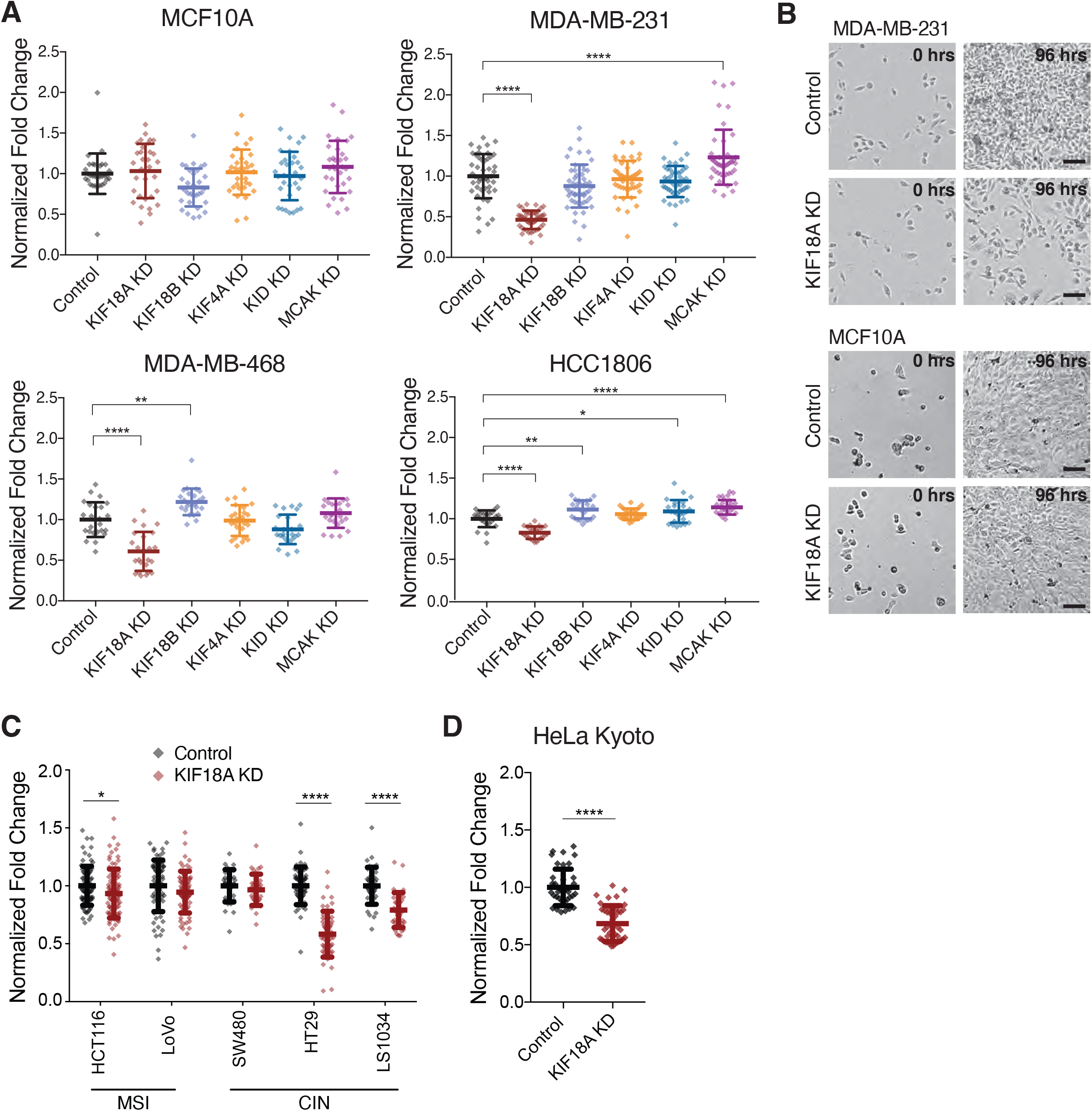
KIF18A is required for the proliferation of chromosomally unstable cells. (A) Fold change in cell density (cells/mm^2^) after 96 hours in the indicated cell lines following knockdown (KD) of kinesin proteins. Data are normalized to cells treated with control siRNA. At least 24 wells per condition from three independent experiments were analyzed. (B) Representative images of MDA-MB-231 and MCF10A cells treated with either control or KIF18A siRNA. Scale bars are 100 microns. (C) Normalized fold change in cell density (cells/mm^2^) of MSI and CIN colorectal cancer cell lines after 96 hours. At least 24 wells per condition from three independent experiments were analyzed in A and C. (D) Normalized fold change in cell density (cells/mm^2^) of HeLa Kyoto cells after 96 hours. Data were generated from two independent experiments. All graphs show mean +/− SD. **** p<0.0001, *** p<0.001, ** p<0.01, * p<0.05

### Loss of KIF18A induces prolonged mitotic delay in CIN tumor cells

KIF18A is required for chromosome alignment in all cells but also promotes spindle assembly checkpoint satisfaction and progression through mitosis in some cell types^17–23^. To determine if proliferation defects seen in KIF18A-depleted CIN cells correlate with KIF18A’s role in promoting timely metaphase-to-anaphase transitions, we compared the effects of KIF18A KD on mitotic progression in CIN cells and near-diploid cells. KIF18A KD led to an increase in the percentage of mitotic CIN cells but did not significantly alter the percentage of mitotic cells within MCF10A or non-CIN CRC cell populations (Fig 2 A-C and Extended Data Fig 4). Quantification of mitotic duration revealed that all cell types displayed a significant increase in the amount of time required to progress from nuclear envelope breakdown (NEB) to anaphase onset (AO) following KIF18A KD (Fig 2 D-F). Consistent with previous work, the magnitude and variance of mitotic delays were larger in KIF18A KD CIN tumor cells than diploid (MCF10A) or near-diploid cells (HCT116) (Fig 2D)^19–21,24,25^. In addition, the cell types most sensitive to KIF18A KD contained a significant subpopulation of cells that failed to complete mitosis during the imaging studies and were arrested for up to 20 hours (Fig 2E). SW480 CIN cells, which were not dependent on KIF18A for proliferation, did not display extended mitotic arrest, suggesting that they are able to compensate for loss of KIF18A in order to complete cell division. These data suggest that proliferation defects in KIF18A-dependent CIN cells may stem from defects that prevent subpopulations of cells from completing mitosis.

**Figure 2.**
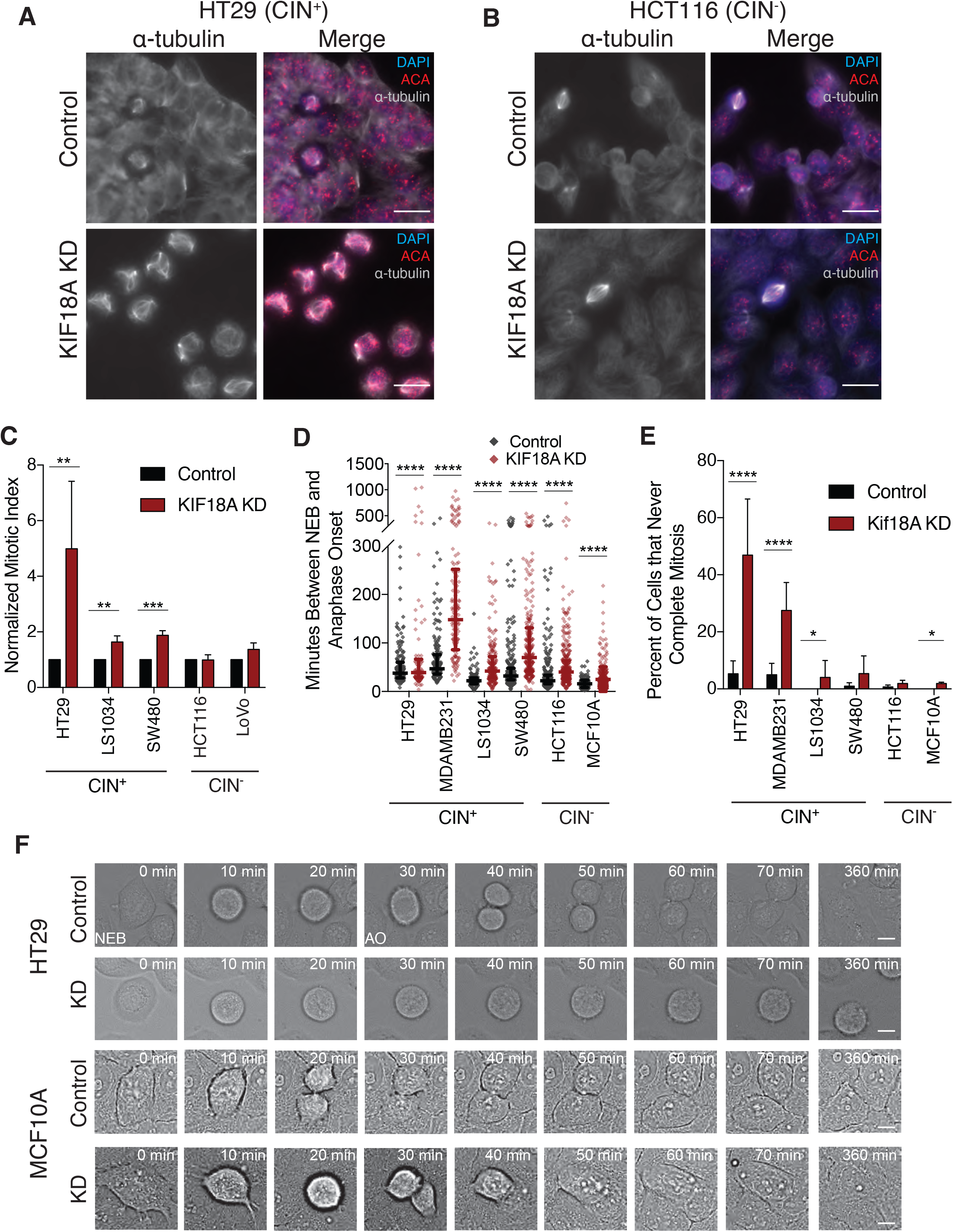
KIF18A depletion causes mitotic arrest in CIN cancer cells. (A,B) Representative images of HT29 (A) or HCT116 cells (B) treated with control or KIF18A siRNAs. Scale bars are 10 microns. (C) Percentage of mitotic cells (mitotic index) observed in fixed populations of control or KIF18A siRNA-treated CRC cells. At least 60 fields from three independent experiments were analyzed per condition. (D) Time between nuclear envelope breakdown (NEB) and anaphase onset (AO) in control or KIF18A siRNA-treated cells. At least 150 cells from three independent experiments were analyzed per condition. (E) Percentage of control or KIF18A siRNA-treated cells that entered mitosis at least 200 minutes before the end of the movie but did not divide. (F) Frames from DIC live cell imaging of HT29 and MCF10A cells treated with control or KIF18A siRNA, showing progression from NEB to AO. Scale bars are 5 microns. All graphs show mean +/− SD. **** p<0.0001, *** p<0.001, ** p<0.01, * p<0.05

### KIF18A-dependent CIN cells form multipolar spindles

Analyses of mitotic spindles in KIF18A KD cells revealed that KIF18A-dependent CIN lines display a significant increase in multipolar spindles compared to non-KIF18A-dependent cell lines (Fig 3 A-B). Multipolar spindles were defined as spindles containing more than two microtubule organizing centers with foci of the pericentriolar component γ-tubulin. Interestingly, the fold-increase in multipolar spindles following KIF18A KD was inversely proportional to the fold-decrease in proliferation for each cell type (Fig 3C). These data indicate that mitotic spindle assembly is abnormal in most KIF18A-dependent CIN cells. However, HeLa cells, which displayed reduced proliferation following KIF18A KD, displayed a slightly different spindle phenotype. A fraction of HeLa cells exhibited fragmented γ-tubulin in response to KIF18A KD but overall did not display a significant increase in multipolar spindles (Fig 3B and Extended Data Fig 5A). These observations suggest that although abnormal γ-tubulin distribution occurs in KIF18A KD HeLa cells, other mechanisms may be keeping the spindles from becoming multipolar. To investigate this further, we examined the effects of KIF18A KD in combination with previously validated siRNAs targeting either CLASP1 or HSET, which have roles in maintaining centrosome integrity and clustering supernumery centrosomes^26–30^. HeLa cells treated with siRNAs against both KIF18A and either HSET or CLASP1 exhibited a significant increase in multipolar spindles with more than two γ-tubulin foci compared to cells treated with siRNAs against a single target (Extended Data Fig 5B). These co-depletion phenotypes were rescued when GFP-KIF18A was inducibly expressed at levels similar to endogenous KIF18A, which were also sufficient to rescue mitotic progression defects caused by KIF18A KD (Extended Data Figure 5B-E). Notably, the combined effects of KIF18A and CLASP1 on spindle morphology were not specific for HeLa cells. Similar results were seen SW480 cells, suggesting this cell type also relies on other mechanisms for centrosome integrity in the absence of KIF18A (Extended Data Fig 5F). Together, these data suggest a function for KIF18A in preserving centrosome integrity that is essential for maintenance of spindle bipolarity in the majority of CIN cells tested.

**Figure 3.**
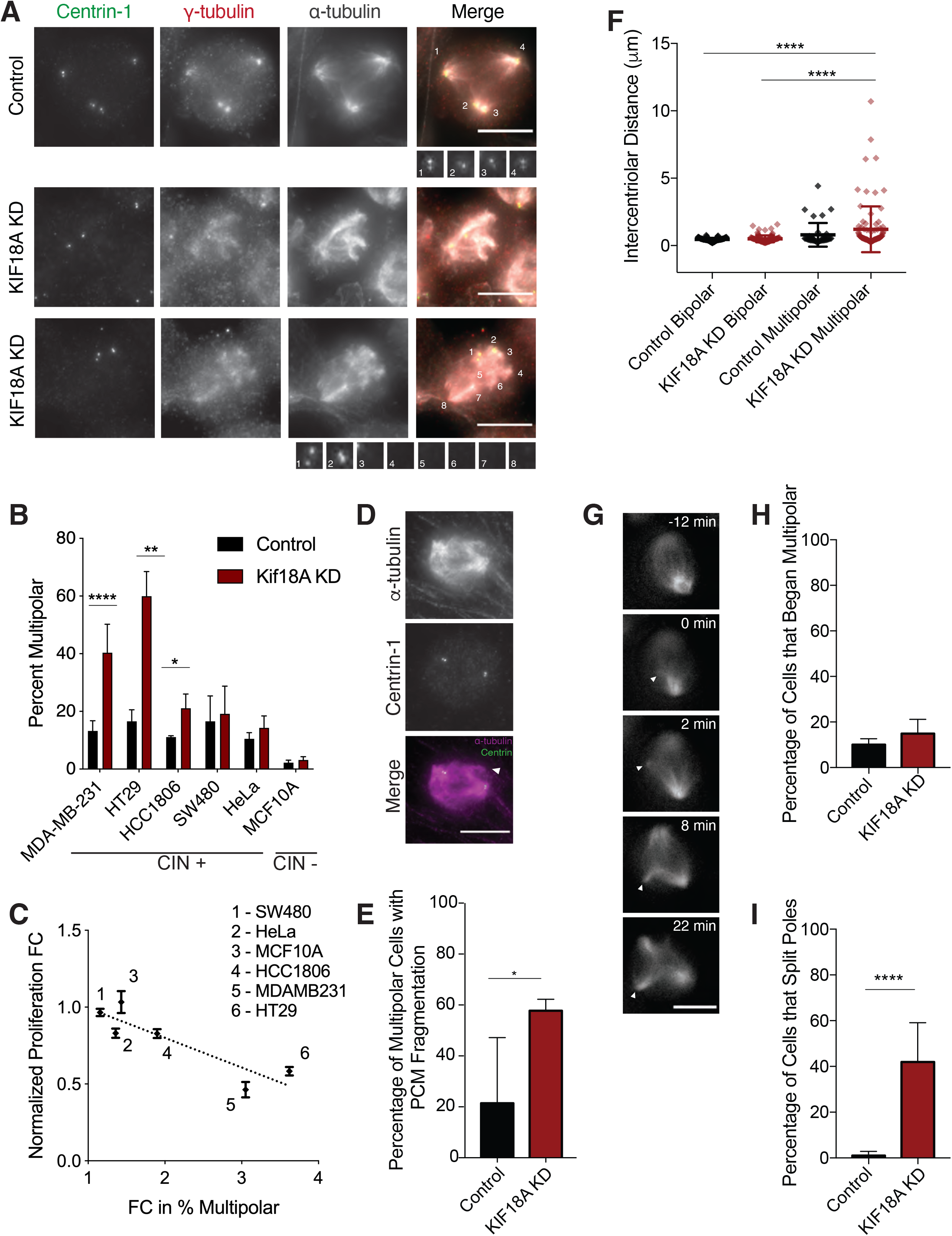
Loss of KIF18A causes centrosome fragmentation in MDA-MB-231 cells. (A) Representative images of MDA-MB-231 cells treated with either control (top) or KIF18A (bottom) siRNA. Pericentriolar material (γ-tubulin) is numbered to show poles with and without centrioles (centrin-1). Scale bars are 10 microns. (B) Percent of mitotic cells with multipolar spindles from fixed cell images of each indicated cell line treated with either control or KIF18A siRNA. Data were generated from at least three independent experiments per cell line. (C) Plot of multipolar spindle percentage as a function of fold-change (FC) in cell number for the indicated cell lines following KIF18A KD. R-squared value is 0.79 using a linear regression model. (D) Representative Images of MDA-MB-231 cell with a third pole lacking centrin-1. Scale bar is 10 microns. (E) Percent of multipolar MDA-MB-231 cells in mitosis with fragmented pericentriolar material (PCM), as indicated by the presence of γ-tubulin puncta lacking centrin-1 puncta. (F) Intercentriolar distance measurements (in microns) for MDA-MB-231 cells in each indicated category. (G) Representative still frames of a live MDA-MB-231 KIF18A KD cell labeled with siR-tubulin. Arrows indicate pole splitting and separation. (H, I) Percent of live, siR-tubulin labeled MDA-MB-231 cells that (H) enter mitosis with more than two spindle poles or (I) split and separate spindle poles during mitosis. Data from three independent experiments. All graphs show mean +/− SD. **** p<0.0001, *** p<0.001, ** p<0.01, * p<0.05

Loss of KIF18A function could lead to multipolar spindles by promoting centrosome amplification, cytokinesis failure, centriole disengagement, or pericentriolar material (PCM) fragmentation^31^. To distinguish among these mechanisms, we analyzed the number and organization of centrioles within multipolar spindles in MDA-MB-231 cells (Fig 3 D-F). The majority of spindles (~75%) in both control and KIF18A KD cells contained four centrioles, indicating that centrosome amplification and cytokinesis failure do not significantly contribute to spindle defects in KIF18A KD cells.

The average distance between paired centrioles was increased in multipolar KIF18A KD cells compared to those in bipolar spindles but was comparable to the average distance measured in multipolar spindles treated with control siRNA (Fig 3F). However, ~60% of multipolar KIF18A KD cells exhibited γ-tubulin containing microtubule organizing centers without centrioles (Fig 3E). A similar trend was observed in HT-29 cells following KIF18A KD (Extended Data Fig 6). Furthermore, live imaging of KIF18A-depleted MDA-MB-231 cells labeled with siR-tubulin revealed an increase in spindle pole fragmentation events but not the number of cells entering mitosis with multiple poles compared to control siRNA treated cells (Fig 3 G-I and Supplementary Videos 1-3). These data suggest that KIF18A KD primarily leads to multipolar spindles by inducing PCM fragmentation.

### KIF18A KD induces multipolar spindles in CIN cells independently of mitotic delay

The fragmentation of centrosomes and formation of multipolar spindles following KIF18A KD could result from abnormal spindle forces caused by altered microtubule dynamics or as a secondary effect of an extended mitotic delay^31^. To determine if a mitotic delay is required for multipolar spindle formation following KIF18A KD, we analyzed spindle morphology in MDA-MB-231 cells depleted of both KIF18A and MAD2, which is required for spindle assembly checkpoint-dependent mitotic arrest^32^. KIF18A/MAD2 KD cells displayed a reduced mitotic index but a similar level of multipolar spindles compared to KIF18A KD cells (Extended Data Fig 7 A-B). Spindle pole splitting in live cells occurred at a range of times after mitotic entry in KIF18A KD cells and at times shortly after NEB in KIF18A/MAD2 KD cells (Extended Data Fig 7 C-E and Supplementary Video 4). The significant decrease in multipolar KIF18A/MAD2 KD cells compared to KIF18A KD alone observed during live imaging may be explained by the limitations inherent to the identification of multipolar spindles in live assays, as poles must split sufficiently far apart to be completely separated in this case. Therefore, the live approach is likely to underestimate the actual time to splitting and percentage of multipolar spindles, especially in cells that exit mitosis quickly. Taken together, these data suggest that loss of KIF18A leads to spindle pole fragmentation in CIN cells and that this defect does not require, but may be enhanced by, a mitotic delay.

### Centrosome fragmentation in KIF18A KD cells requires dynamic microtubules

KIF18A functions to suppress microtubule growth within mitotic spindles^18,33^, suggesting that abnormal microtubule dynamics in KIF18A KD cells may contribute to centrosome fragmentation. We tested this idea by reducing microtubule polymerization or depolymerizing microtubules completely via treatment of KIF18A KD MDA-MB-231 cells with 20 nM paclitaxel or 5 μM nocodazole, respectively^34,35^. KIF18A KD cells treated with either paclitaxel or nocodazole for 3 hours before fixation displayed significantly fewer multipolar spindles than KIF18A KD cells treated with DMSO (Fig 4A). These data indicate that dynamic microtubules are required for KIF18A KD-induced centrosome fragmentation.

**Figure 4.**
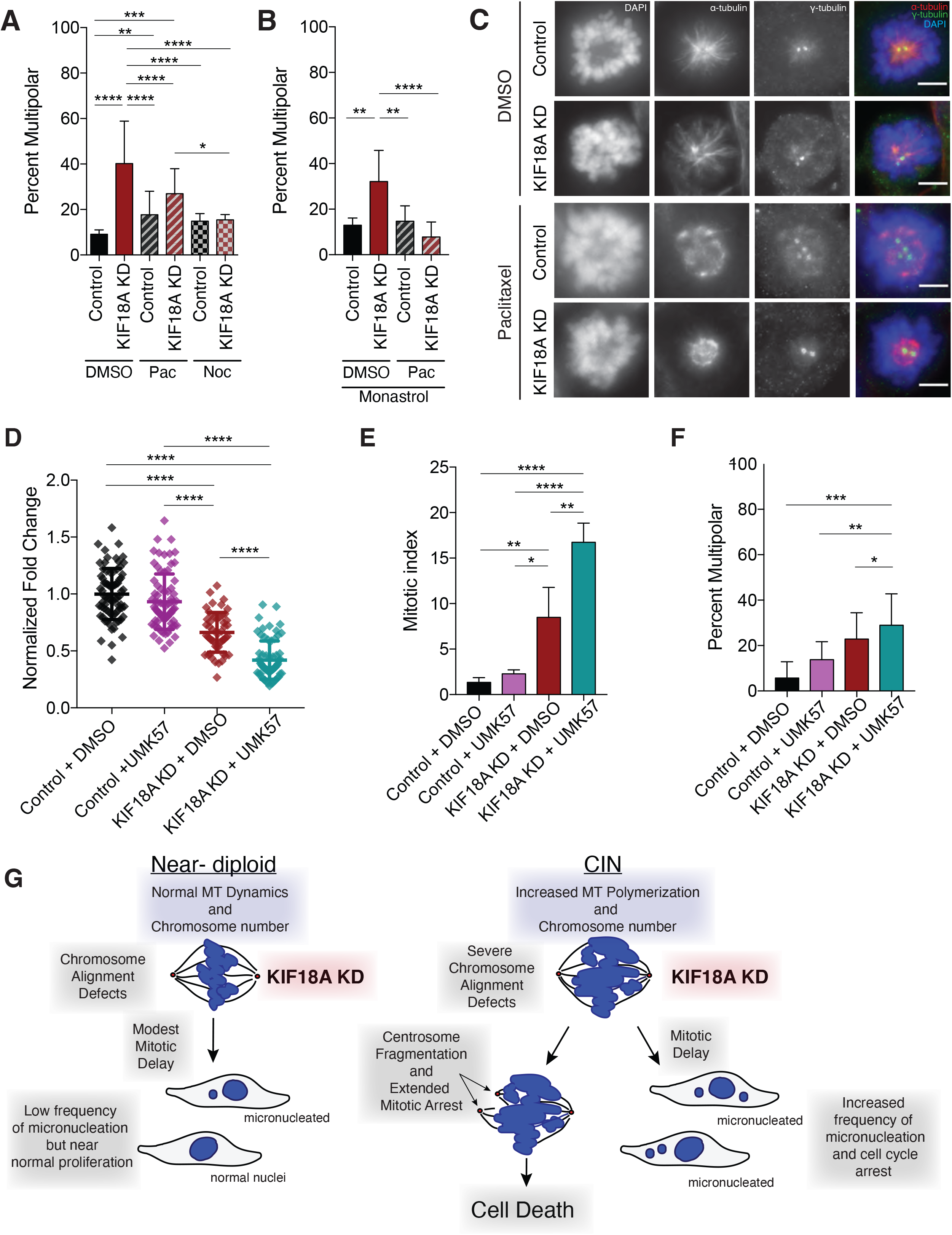
KIF18A KD-induced defects depend on dynamic microtubules and are enhanced by increased KIF2C/MCAK activity. (A) Percent of MDA-MB-231 cells with multipolar spindles in control or KIF18A KD cells treated with either DMSO, 20 nM Paclitaxel (Pac), or 5 μM Nocodazole (Noc) for three hours. At least 150 mitotic cells were analyzed per condition from three independent experiments. (B) Percent of monopolar MDA-MB-231 cells with three or more γ-tubulin puncta in control or KIF18A KD cells treated with both monastrol (20 μM) and either DMSO or 20 nM Paclitaxel. At least 100 monopolar cells were analyzed per condition from three independent experiments. (C) Representative images of MDA-MB-231 cells treated with 20 μM monastrol and either DMSO or 20 nM Paclitaxel. Scale bar is 5 microns. (D) Fold change in cell density after 96 hours in MDA-MB-231 cells treated with the specified siRNAs and either 500 nM UMK57 or DMSO. At least 52 wells per condition from three independent experiments were analyzed. (E,F) Percent of total mitotic cells (E) and mitotic cells with multipolar spindles (F) in fixed populations after the indicated treatment. At least 60 fields from three independent experiments were analyzed per condition. All graphs show mean +/− SD. **** p<0.0001, *** p<0.001, ** p<0.01, * p<0.05 (G) Schematic model for selective dependence of CIN cells on KIF18A function. See Discussion text for details.

Paclitaxel treatment causes an increase in multipolar spindles in both somatic diploid cells and aneuploid cancer cells^13,36,37^, whereas KIF18A KD preferentially affected spindle morphology in the CIN cells tested here (Fig 3A-B). To directly compare the two treatments, we analyzed MDA-MB-231 and MCF10A cells after KIF18A KD or incubation with a clinically relevant dose of paclitaxel (10 nM) for 24 hours in parallel^13^. Paclitaxel treatment led to increased multipolar spindles in both MDA-MB-231 cells and MCF10A cells, while KIF18A KD only induced multipolar spindles in MDA-MB-231 cells (Extended Data Fig 8). Thus, paclitaxel and KIF18A KD induce a similar spindle phenotype but appear to act via antagonistic mechanisms, consistent with their opposite effects on spindle microtubule dynamics. Furthermore, sensitivity to KIF18A loss of function is higher in the CIN cell lines tested here than in near-diploid MCF10A cells.

### Centrosome fragmentation in KIF18A KD cells does not require bipolar spindle formation

Our live imaging studies indicated that KIF18A KD MDA-MB-231 and HT29 cells form bipolar spindles before centrosome fragmentation occurs (Fig 3, Extended Data Fig 6, and Supplementary Videos 1-3). Therefore, altered microtubule dynamics in KIF18A KD cells could lead to centrosome fragmentation by disrupting the balance of pushing and pulling forces within bipolar spindles. To test this idea, we assayed the number of γ-tubulin foci in MDA-MB-231 cells treated with the KIF11 inhibitor monastrol, which induces monopolar spindles by preventing KIF11-dependent anti-parallel microtubule sliding forces^38^. We found that centrosome fragmentation still occurred in monopolar KIF18A KD cells and could be reduced by co-treatment with paclitaxel (Fig 4 B-C). Live imaging of monastrol treated cells expressing RFP-pericentrin to label centrosomes revealed that centrosomes begin intact in monopolar KIF18A KD cells and subsequently fragment (Supplementary Video 5). These data suggest that neither bipolar spindles nor the forces generated via KIF11-dependent microtubule sliding are required for centrosome fragmentation in the absence of KIF18A.

### The effects of KIF18A KD are enhanced by activation of MCAK/KIF2C

In addition to suppressing the growth of kinetochore microtubules, KIF18A is required to decrease kinetochore microtubule turnover^18,39^. Thus, we reasoned that increased kinetochore microtubule turnover may contribute to the prolonged mitotic delays and destabilized spindles observed in KIF18A KD CIN cells. To test this, we treated cells with a small molecule (UMK57) that promotes kinetochore microtubule turnover by increasing the activity of the depolymerizing kinesin MCAK/KIF2C^40^. Treatment of KIF18A-depleted MDA-MB-231 cells with UMK57 (500 nM) decreased proliferation and significantly increased the mitotic index compared to KIF18A KD cells treated with DMSO (Fig 4 D-E and Extended Data Fig 9A). The same concentration of UMK57 had no impact on the proliferation of control siRNA-treated cells (Fig 4D). UMK57 treatment of KIF18A KD cells also led to a small but significant increase in multipolar spindles, and this effect was replicated in cells with increased global MCAK/KIF2C activity, due to over-expression of mCherry-MCAK, or increased MCAK/KIF2C activity at centromeres, due to expression of mCherry-CPB-MCAK^41^ (Fig 4F and Extended Data Fig 9B). Furthermore, in live cells labeled with siR-tubulin, co-depletion of both KIF18A and KIF2C reduced multipolar spindle formation compared to depletion of KIF18A alone, while KIF18A KD cells treated with UMK57 displayed increased spindle pole fragmentation (Extended Data Fig 9 C-F and Supplementary Video 6). These data indicate that loss of KIF18A function combined with increased KIF2C function, particularly at centromeres, synergistically disrupts mitotic progression and spindle bipolarity.

## Discussion

Our data support a model in which the altered microtubule dynamics in mitotic CIN cells make them particularly dependent on KIF18A to reduce kinetochore microtubule turnover and limit microtubule growth. In the absence of KIF18A activity, centrosome fragmentation occurs and mitotic progression is slowed or prevented. Importantly, these effects were not observed in the near-diploid, chromosomally stable cells tested here, which is consistent with previous observations that reduction of KIF18A activity leads to longer spindle assembly checkpoint-dependent delays in cancer cells than diploid somatic cells^17,19–23^. KIF18A is also largely dispensable for proliferation of diploid somatic cells *in vivo* but is necessary for tumor growth. *Kif18a* mutant mice display an early growth delay and germline development defects^19,42^. However, the growth of both induced CRC and xenografted TNBC tumors in mouse models are dependent on KIF18A^43,44^. Thus, KIF18A may be an effective target to specifically inhibit the growth of CIN tumor cells, while inducing relatively low toxicity in somatic, diploid cells.

These data raise the important question of why CIN cells would depend more on KIF18A for successful mitosis than chromosomally stable cells. CIN cells exhibit increased rates of spindle microtubule polymerization and altered turnover of kinetochore microtubules^2,3^, which may confer an enhanced dependence on KIF18A’s function to suppress microtubule growth^18,45,46^. Our results suggest that in the absence of KIF18A activity, maintenance of kinetochore microtubule attachments and centrosome integrity are compromised in CIN cells, subsequently leading to extended mitotic arrest and centrosome fragmentation. These phenotypes were suppressed by treatments that reduce microtubule dynamics (nocodazole, paclitaxel, and KIF2C KD) and were enhanced by treatments that increase microtubule dynamics (UMK57, KIF2C overexpression) or compromise centrosome integrity (CLASP1 or HSET KD). Taken together, these data support a model in which the combined effects of increased microtubule polymerization in CIN cells and loss of KIF18A’s microtubule growth suppression create a force imbalance within spindles that reduces centrosome integrity (Fig 4G). Additionally, this force imbalance does not require a bipolar spindle configuration or KIF11-dependent microtubule sliding within the spindle, as indicated by centrosome fragmentation in monastrol treated cells.

We also observed that a significant fraction of CIN cells are able to complete mitosis after a mitotic delay. Loss of KIF18A could also slow the growth of this population by inducing chromosome segregation errors. KIF18A KD increases the frequency of micronucleus formation as a result of chromosome alignment defects, and these micronucleated cells display reduced proliferation^20^. In support of this idea, recent work indicates that the frequency of KIF18A KD-dependent micronuclei is increased in cells with elevated chromosome number and correlates with reduced proliferation in aneuploid cells^47,48^.

The mitotic defects induced by KIF18A KD, multipolar spindles and micronuclei, are similar to those proposed to underlie the anti-tumor activity of paclitaxel^11,14^, yet KIF18A KD and paclitaxel have antagonistic effects on microtubule dynamics and lead to reduced multipolar spindle formation when combined. This suggests that increases or decreases in microtubule dynamics may compromise spindle integrity in CIN tumor cells, an idea supported by observations that multipolar spindles form in cells with dampened microtubule dynamics due to KIF18A overexpression^45^. Our tests of other kinesins that control spindle microtubule dynamics and chromosome movements suggest the observed dependence of CIN cells on KIF18A is unique. In agreement, two recent, large-scale bioinformatics studies identified *Kif18A*, but not other kinesins, as a gene specifically required for the growth of cells displaying aneuploidy or whole genome duplication^47,48^. Thus, KIF18A represents a potential target for exploiting vulnerabilities specific to a significant fraction of tumor cells displaying CIN or aneuploidy.

---

**Extended Data Figure 1.**
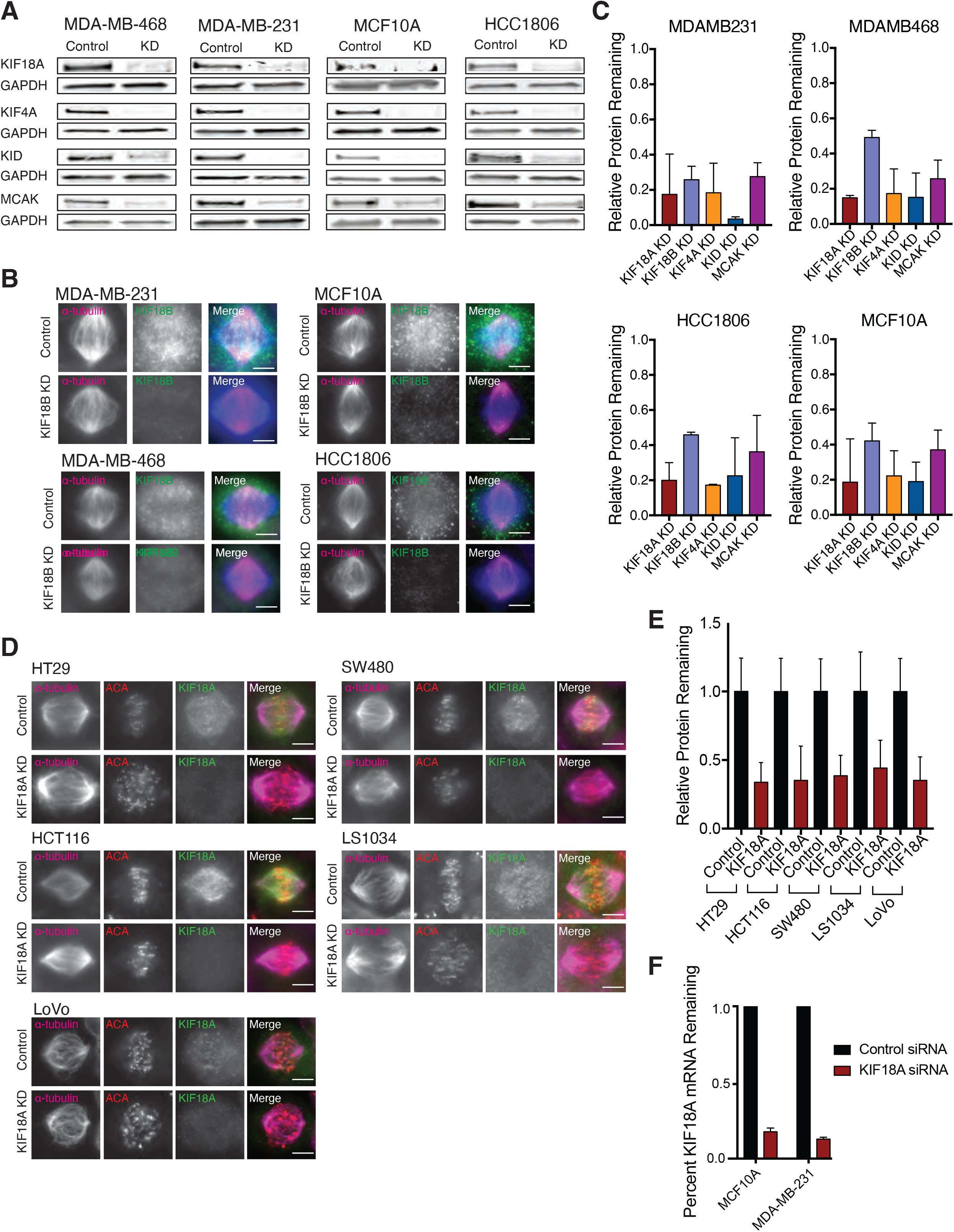
Kinesins are effectively depleted by siRNA in breast and colorectal cell lines. (A) Western blots showing siRNA knockdown (KD) efficiencies for the indicated kinesins in TNBC and diploid breast epithelial cells. (B) Immunofluorescence images demonstrating efficiency of KIF18B KD in TNBC and diploid breast epithelial cells. Scale bar is 10 microns. (C) Quantification of kinesin knockdowns in TNBC and diploid breast epithelial cells from 2-3 independent replicates. Relative remaining protein indicates the proportion of each kinesin remaining in cells after siRNA knockdown (measured via western blot or immunofluorescence) relative to control. (D) Immunofluorescence images demonstrating efficiency of KIF18A siRNA-mediated knockdown in CRC cell lines. Scale bar is 10 microns. (E) Quantification of kinesin knockdowns in CRC cell lines from 2-3 independent replicates. Relative remaining protein was measured via immunofluorescence, and all values within each cell line were normalized to control. (F) Quantitative PCR measurements of KIF18A mRNA levels after siRNA-mediated knockdown in diploid breast epithelial cells and one TNBC cell line. Data from two independent replicates. All graphs show mean +/− SD.

**Extended Data Figure 2.**
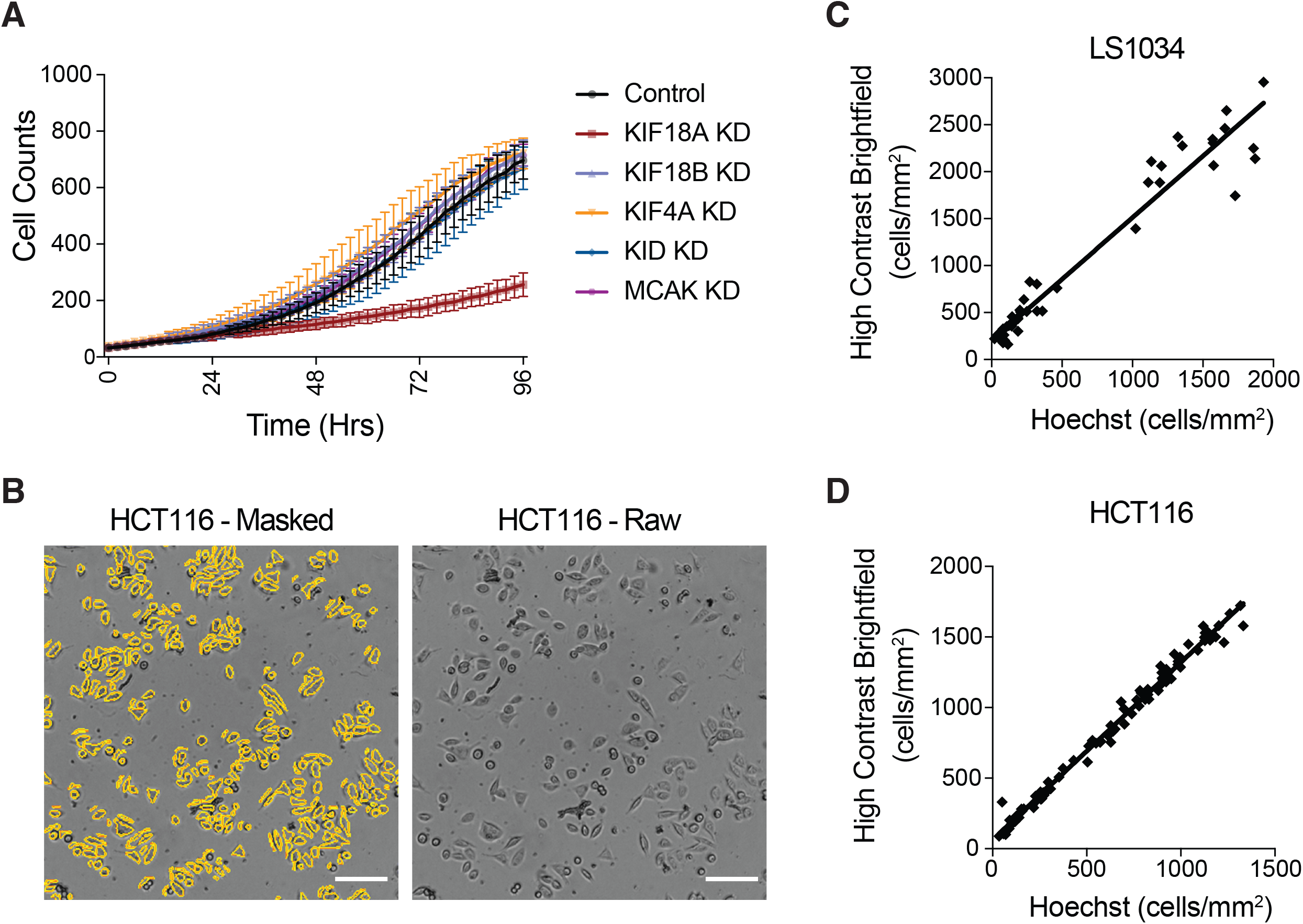
Kinetic cell proliferation assay validation. (A) Example trace of MDA-MD-231 cell density (cells/mm^2^) as a function of time over 96 hours. (B) Representative images of HCT116 cells showing the masks created for automated cell counting. (C-D) Scatterplots of automated (C) LS1034 and (D) HCT116 cell counts using high-contrast brightfield microscopy as a function of cell counts of the same fields using a nuclear dye (Hoechst). Linear correlation indicates consistency in automated cell counting across different cell densities.

**Extended Data Figure 3.**
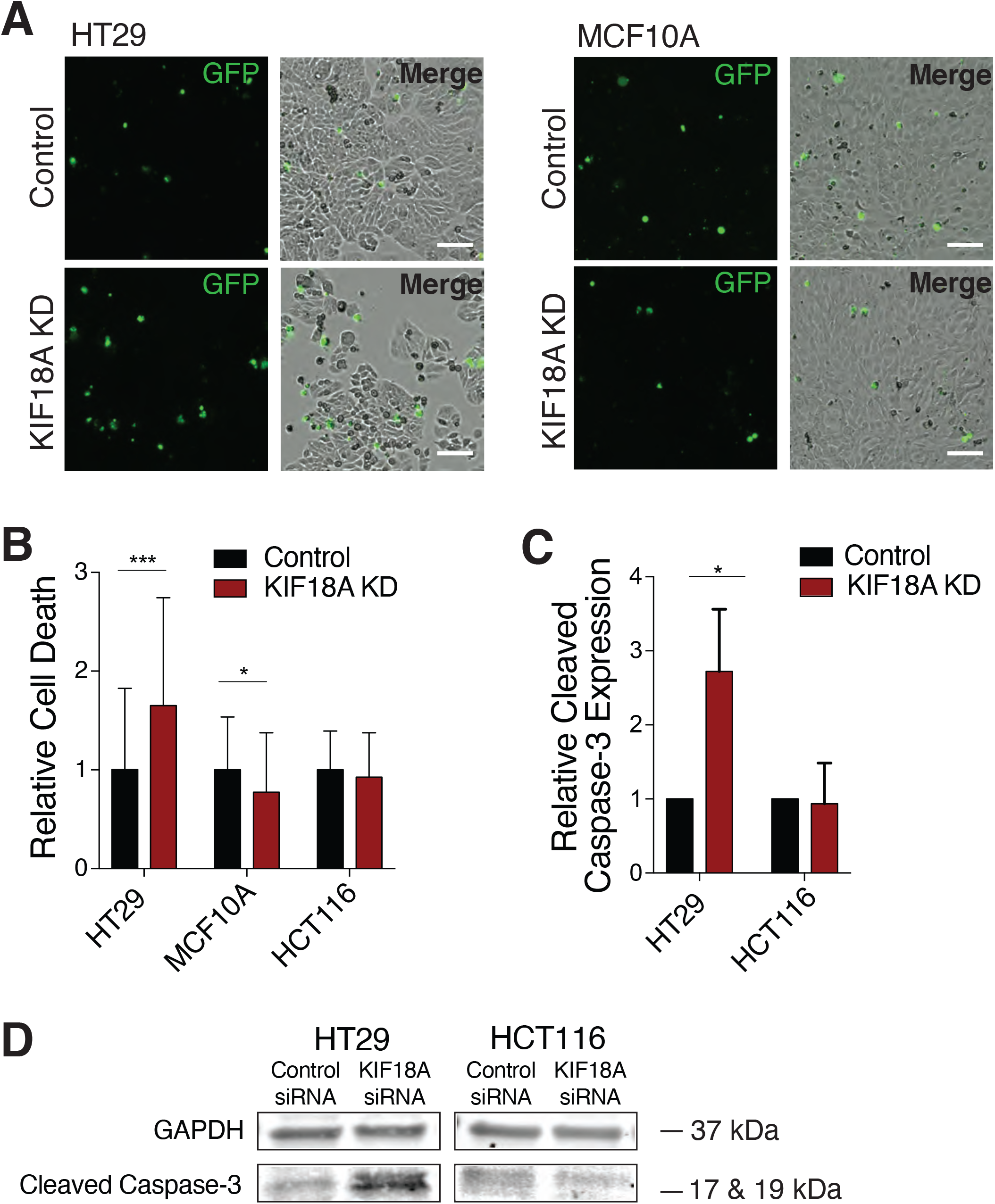
KIF18A depletion increases cell death in CIN cells. (A) Representative images of HT29 and MCF10A cells labeled with Celltox Green cytotoxicity dye five days after siRNA transfection. Scale bars are 100 microns. (B) Relative cell death calculated as the normalized ratio of the change in Celltox-stained cell density to the change in total cell density over 96 hours. A total of at least 68 wells from three independent experiments were analyzed. (C) Relative expression of cleaved-caspase 3 measured via Western blot for each condition. Results are from three independent experiments. (D) Western blot showing representative cleaved-caspase 3 (CC3) expression levels. All graphs show mean +/− SD. **** p<0.0001, *** p<0.001, ** p<0.01, * p<0.05

**Extended Data Figure 4.**
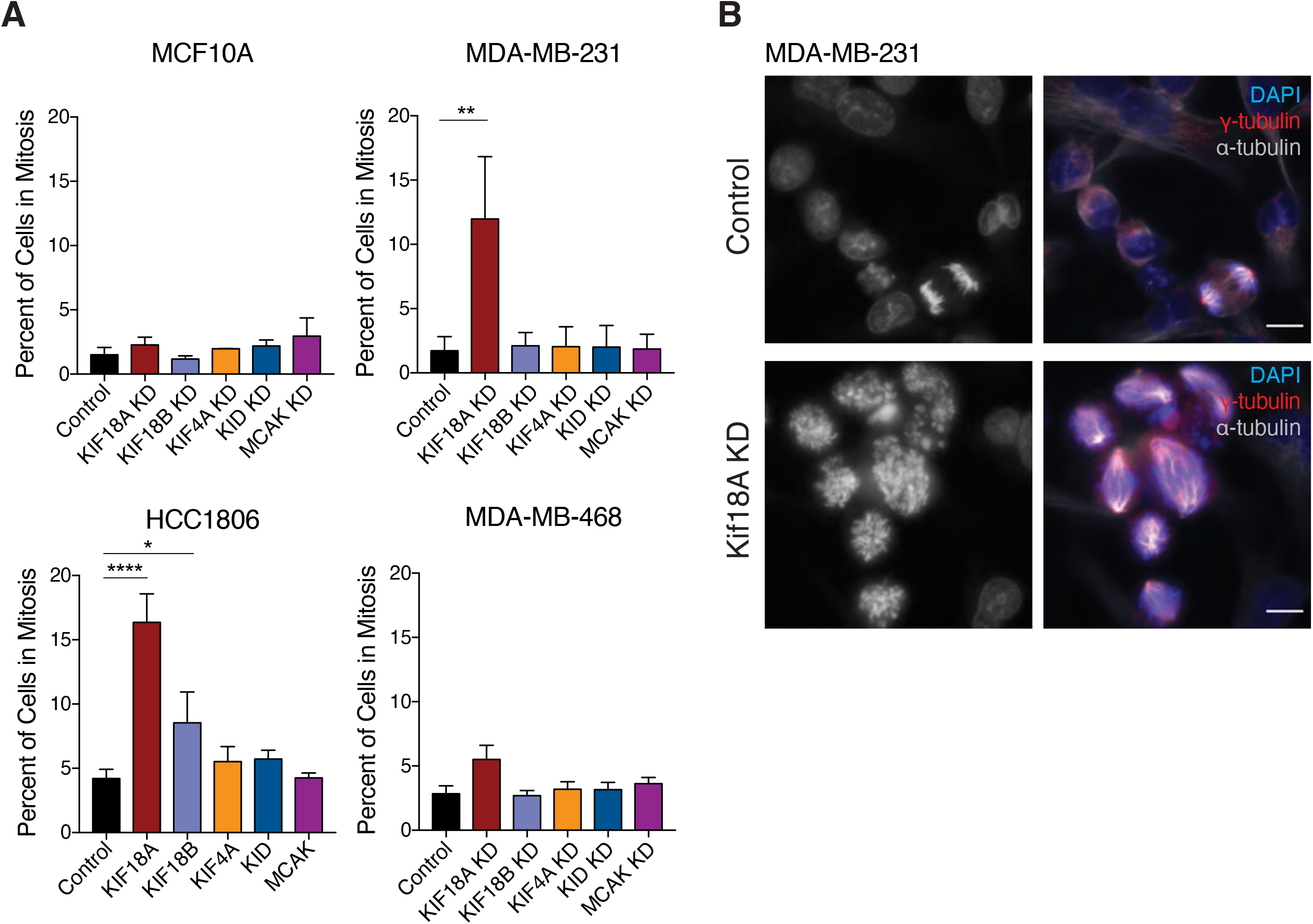
KIF18A KD increases the percentage of cells in mitosis for TNBC cells, but not for diploid breast epithelial cells. (A) Percent of cells in mitosis, as determined from fixed cell images, 48 hours after siRNA-mediated knockdown (KD) of the specified kinesins. Results are from three independent experiments. (B) Representative images of MDA-MB-231 cells treated with either control or KIF18A siRNA. Scale bar is 10 microns. All graphs show mean +/− SD. **** p<0.0001, *** p<0.001, ** p<0.01, * p<0.05

**Extended Data Figure 5.**
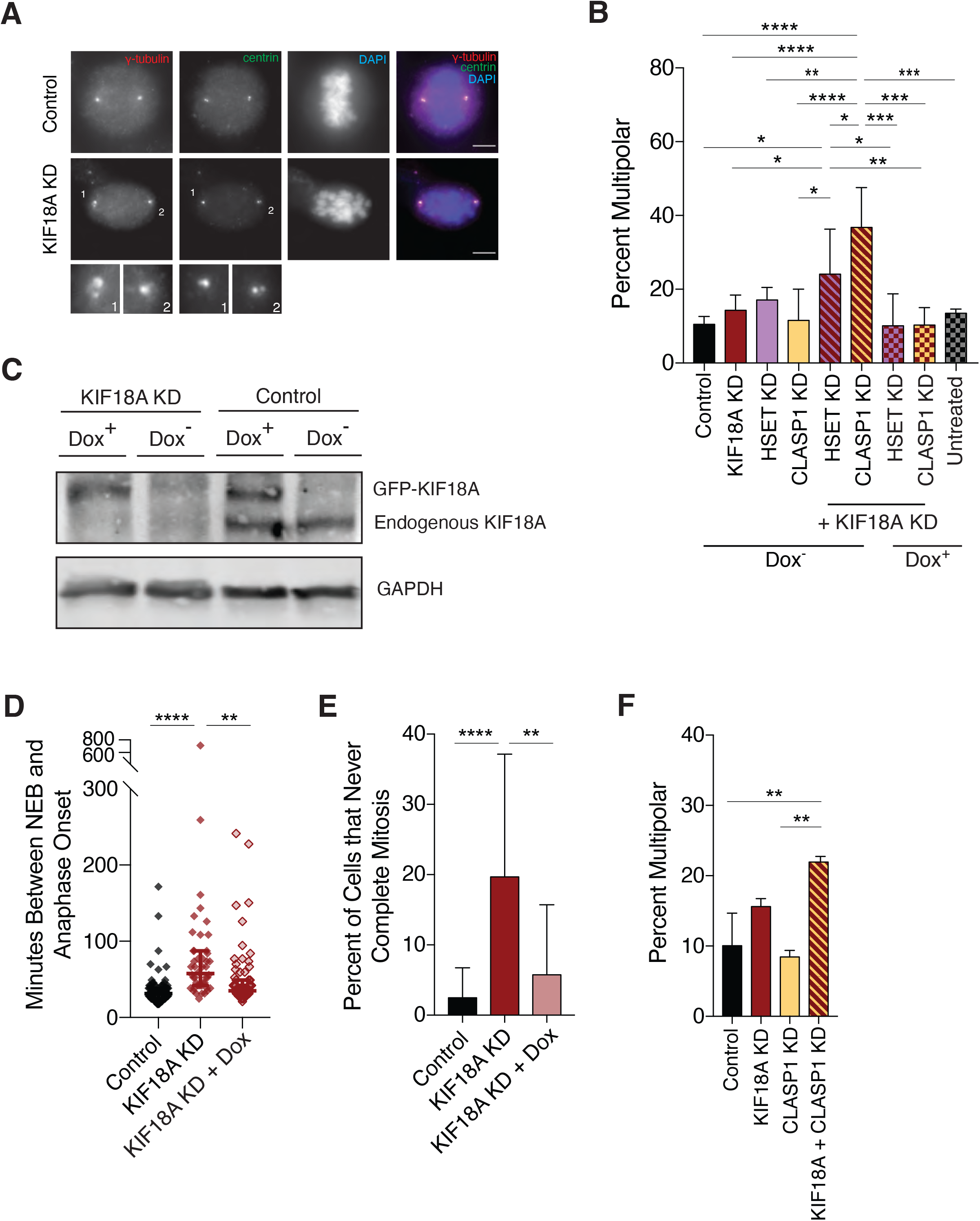
Co-depletion of KIF18A and CLASP1 or HSET lead to synergistic increases in multipolar spindle formation. (A) Representative images of HeLa Kyoto cells treated with either control or KIF18A siRNA. Insets show enlarged images of the numbered poles in the KIF18A KD cell. Scale bar is 10 microns. (B) Percent of HeLa Kyoto cells with multipolar spindles in each of the indicated single or double knockdowns, with or without induction of GFP-KIF18A via doxycycline. Results are from three independent experiments. (C) Western blot depicting the amount of either endogenous KIF18A or GFP-KIF18A in cells treated with KIF18A or control siRNA with or without the addition of doxycycline. (D) Time between nuclear envelope breakdown (NEB) and anaphase onset for HeLa Kyoto cells treated with control or KIF18A siRNA with or without the addition of doxycycline. Results from three independent experiments. (E) Percent of HeLa Kyoto cells that fail to complete mitosis after treated with either control or KIF18A siRNA with or without doxycycline. Results from three independent experiments. (F) Percent of SW480 cells with multipolar spindles in each of the indicated single or double knockdowns. Results from two independent experiments. All graphs show mean +/− SD. **** p<0.0001, *** p<0.001, ** p<0.01, * p<0.05

**Extended Data Figure 6.**
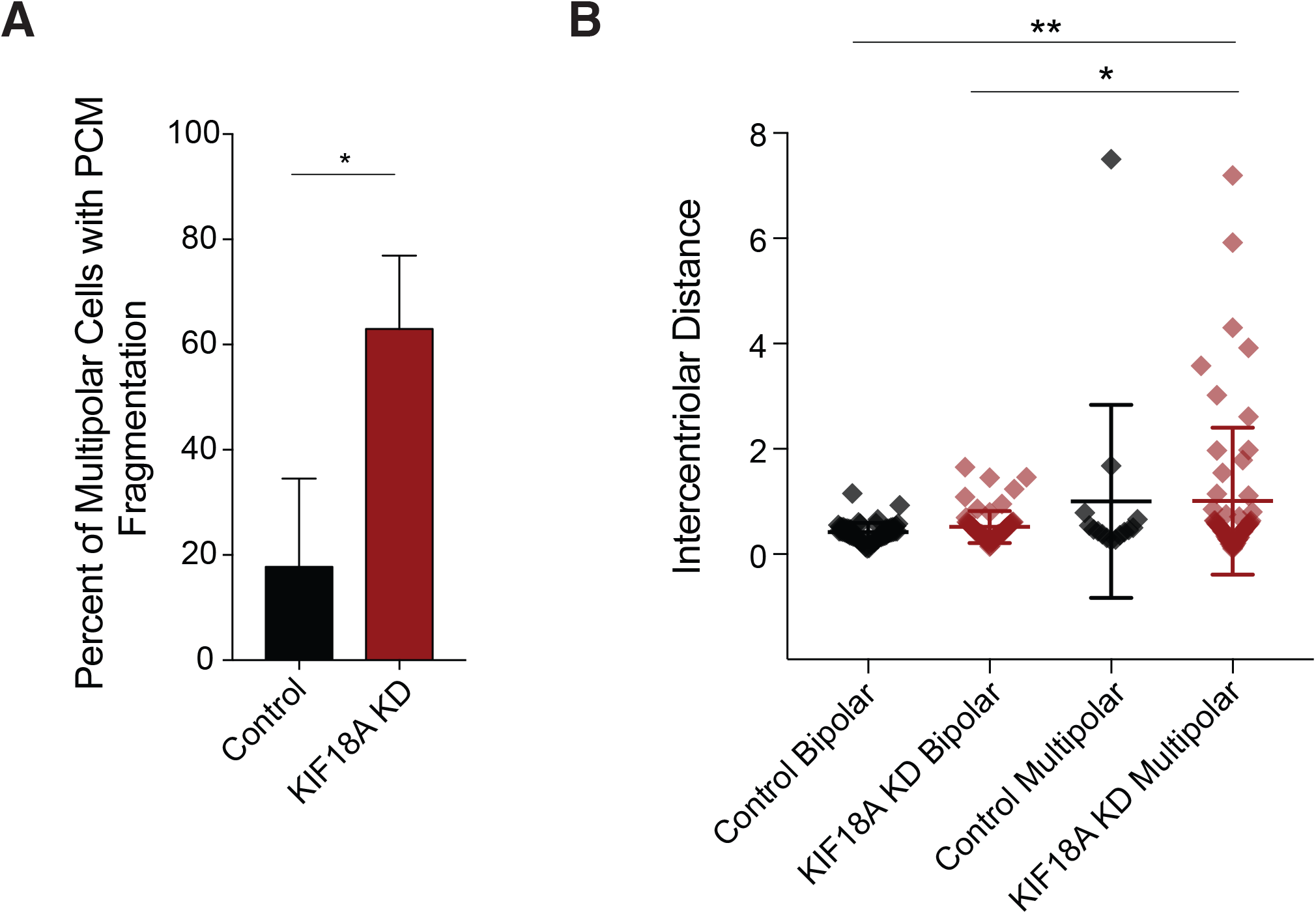
Loss of KIF18A causes centrosome fragmentation in HT29 cells. A. Percent of HT29 cells with fragmented pericentriolar material (PCM), as indicated by the presence of γ-tubulin puncta lacking centrin-1. Results from three independent experiments. B. Intercentriolar distance measurements (in microns) for HT29 cells in each indicated category. Between 15-63 measurements were made per category. All graphs show mean +/− SD. **** p<0.0001, *** p<0.001, ** p<0.01, * p<0.05

**Extended Data Figure 7.**
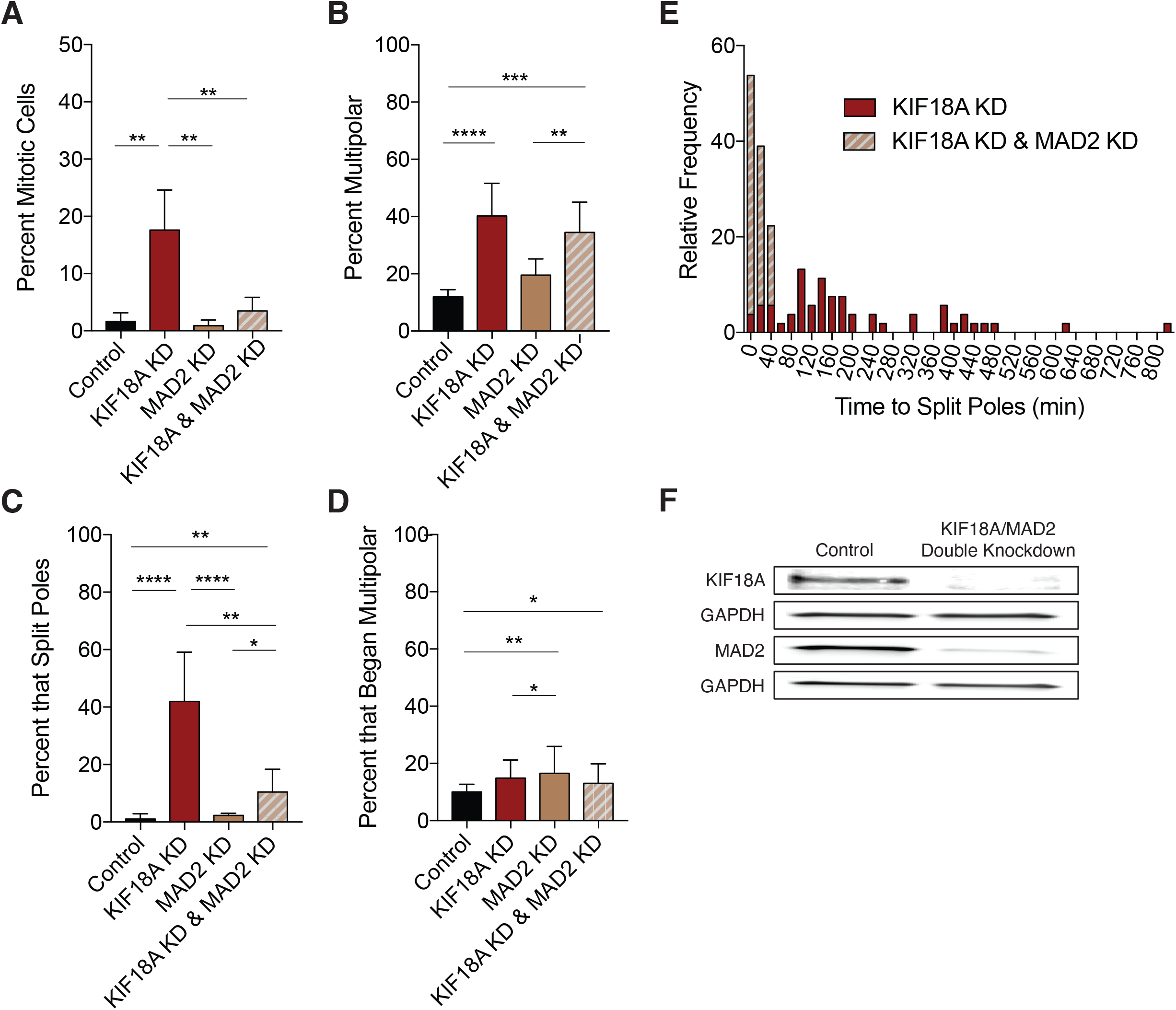
Spindle checkpoint inhibition rescues mitotic arrest but not multipolar spindle formation caused by KIF18A KD. (A-B) Percent of fixed MDA-MB-231 cells (A) in mitosis or (B) with multipolar spindles after the indicated siRNA KD. Results are from three independent experiments. (C-D) Percent of live, siR-tubulin labeled MDA-MD-231 cells that (C) split poles during mitosis or (D) entered mitosis with more than two spindle poles. Results are from two independent experiments. (E) Stacked histogram showing relative frequencies of the duration of time between NEB and pole splitting for siR-tubulin labeled MDA-MB-231 cells following KIF18A KD or KIF18A/MAD2 KD. (F) Western blots depicting the amount of each specified protein remaining after treatment with either a double dose of control siRNA or a combination of KIF18A and MAD2 siRNA. All graphs show mean +/− SD. **** p<0.0001, *** p<0.001, ** p<0.01, * p<0.05

**Extended Data Figure 8.**
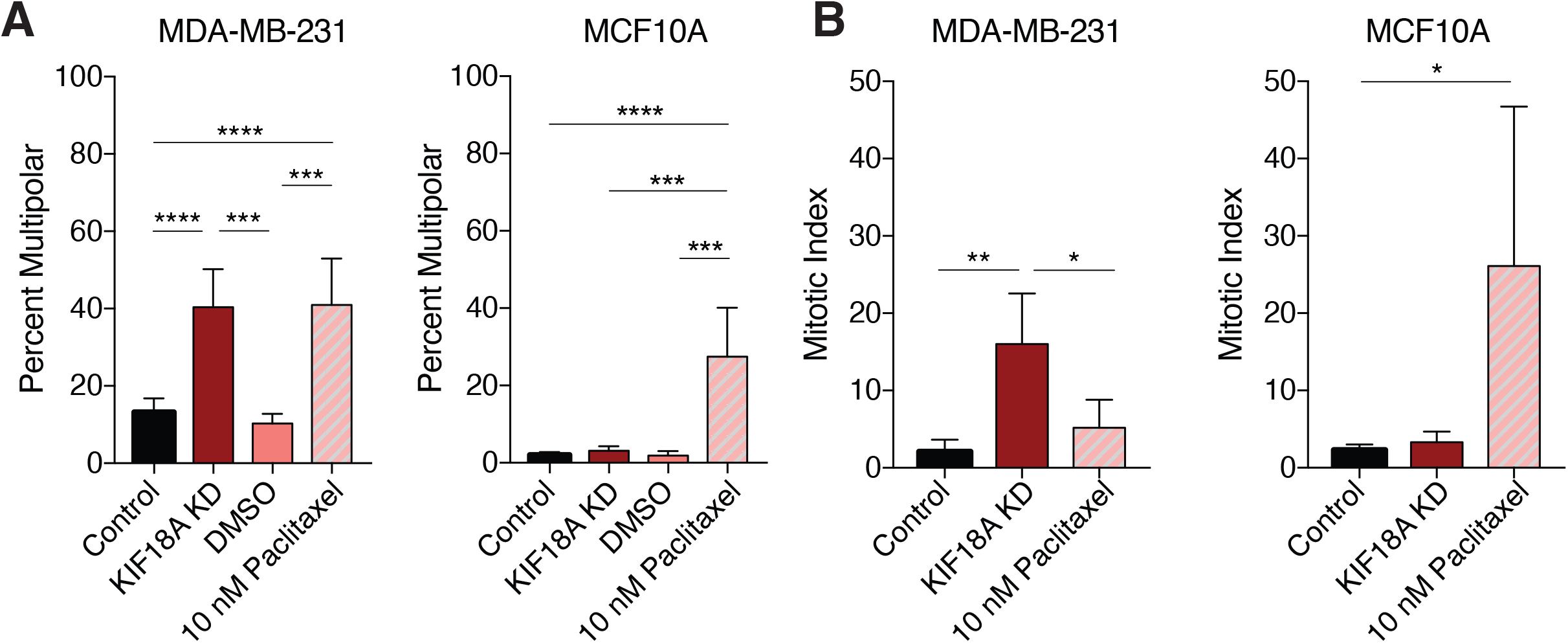
KIF18A KD and paclitaxel treatment cause similar mitotic defects in CIN MDA-MB-231 cells, but not diploid MCF10A cells. (A) Percentage of mitotic cells with multipolar spindles in fixed MDA-MB-231 or MCF10A cells treated with control siRNAs, KIF18A siRNAs, 10 nM paclitaxel, or DMSO. (B) Percentage of fixed MDA-MB-231 and MCF10A cells in mitosis following treatment with control siRNAs, KIF18A siRNAs, 10 nM paclitaxel, or DMSO. All graphs show mean +/− SD from 3 independent experiments. **** p<0.0001, *** p<0.001, ** p<0.01, * p<0.05

**Extended Data Figure 9.**
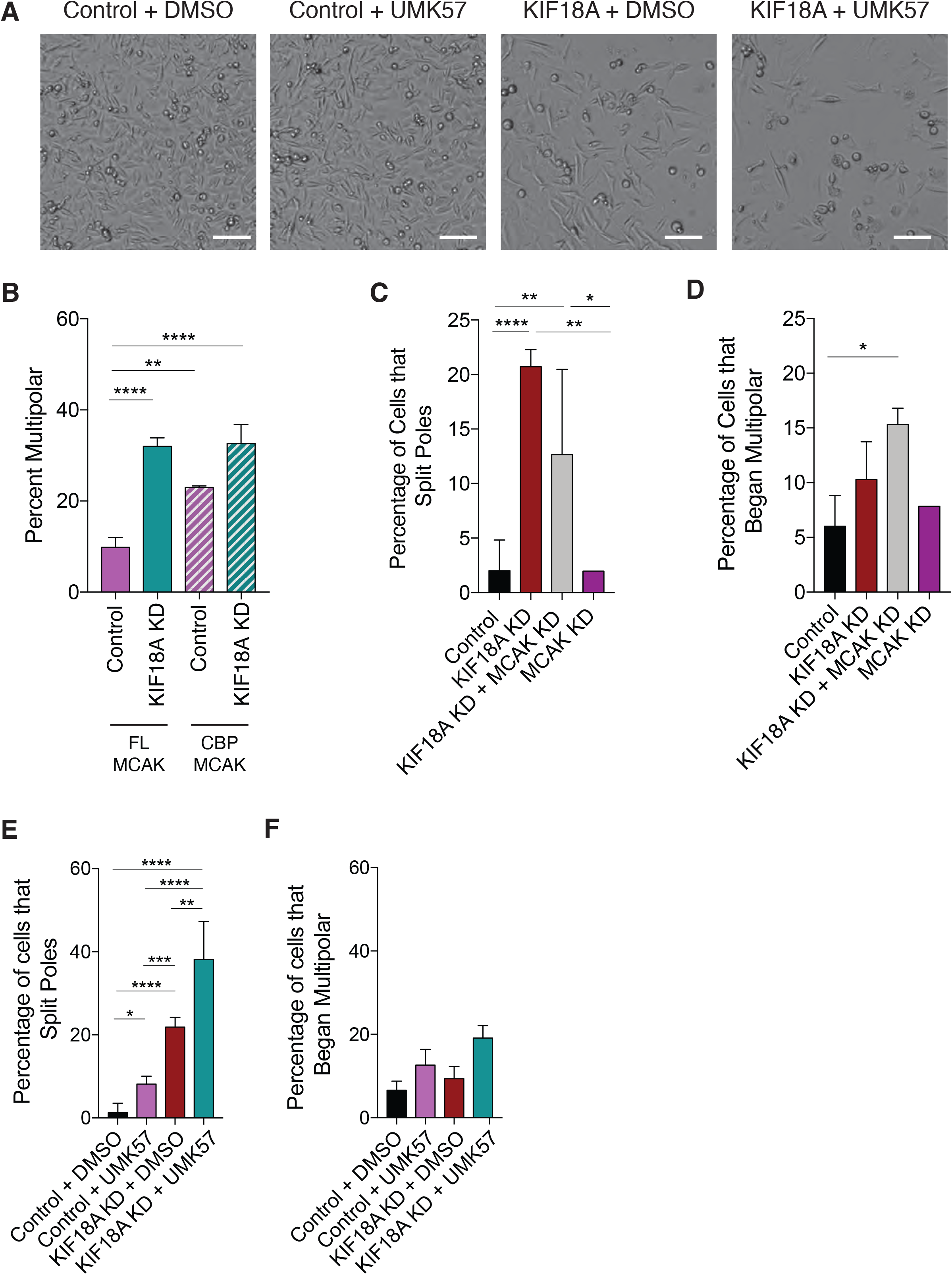
Proliferation and multipolar spindle defects caused by KIF18A KD are sensitive to changes in KIF2C/MCAK activity. (A) Representative images of MDA-MB-231 cell density 96 hours after the start of high-contrast brightfield imaging. Cells were treated with either control or KIF18A siRNA in combination with DMSO or 500 nM UMK57. Scale bar is 100 microns. (B) Percent of MDA-MB-231 cells with multipolar spindles in cells treated with the indicated siRNAs and transfected with mCh-MCAK-FL or mCh-CPB-MCAK. Results are from two independent experiments. (C-D) Percent of live, siR-tubulin labeled MDA-MB-231 cells that (C) split poles or (D) entered mitosis with more than two spindle poles after the indicated single or double knockdowns. MCAK KD results are from one experiment; all other results are from two independent experiments. (E-F) Percent of live, siR-tubulin labeled MDA-MB-231 cells that (E) split poles or (F) entered mitosis with more than two spindle poles after the indicated treatments. Results from three independent experiments. (D) Representative immunofluorescence images of mitotic MDA-MB-231 cells treated with either control or KIF18A siRNA in combination with either DMSO or 500 nM UMK57. Scale bars are 10 microns.

## Supplementary Videos

**Supplementary Video 1.** Representative time-lapse movie of bipolar division in a siR-tubulin labeled MDA-MB-231 cell treated with control siRNA. Images were acquired every 2 min and are played back at 7 frames per second.

**Supplementary Video 2.** Representative time-lapse movie of spindle pole splitting in a siR-tubulin labeled MDA-MB-231 cell treated with KIF18A siRNA. Images were acquired every 2 min and are played back at 7 frames per second.

**Supplementary Video 3.** Representative time-lapse movie of a siR-tubulin labeled MDA-MB-231 cell treated with KIF18A siRNA entering mitosis with multiple spindle poles. Images were acquired every 2 min and are played back at 7 frames per second.

**Supplementary Video 4.** Representative time-lapse movie of spindle pole splitting in a siR-tubulin labeled MDA-MB-231 cell treated with KIF18A and MAD2 siRNAs. Images were acquired every 2 min and are played back at 7 frames per second.

**Supplementary Video 5.** Representative time-lapse movie of centrosome fragmentation in a monopolar MDA-MB-231 cell expressing mRFP-pericentrin after treatment with KIF18A siRNA and 20 μM monastrol. Images were acquired every 2 min and are played back at 7 frames per second.

**Supplementary Video 6.** Representative time-lapse movie of spindle pole splitting in a siR-tubulin labeled MDA-MB-231 cell treated with KIF18A siRNA and 500 nM UMK57. Images were acquired every 2 min and are played back at 7 frames per second.

## Methods

### Cell Culture and Transfections

HT29, LoVo, SW480, LS1034, HCC1806, HCT116, MCF10A, MDA-MB-231, and MDA-MB-468 cells were purchased from ATCC. HeLa Kyoto acceptor cells for recombination mediated cassette exchange were previously described. All cell lines were validated by STR DNA fingerprinting using the Promega GenePrint® 10 System according to manufacturer’s instructions (Promega #B9510). HT29, LoVo, SW480, MDA-MB-231, and MDA-MB-468 cells were cultured in DMEM/F-12 medium (Gibco) supplemented with 10% FBS (Gibco) and 1% penicillin/streptomycin (pen/strep). LS1034 and HCC1806 cells were cultured in RPMI 1640 medium (Gibco) with 10% FBS and 1% pen/strep. HCT116 cells were cultured in McCoys 5A media (Gibco) with 10% FBS and 1% pen/strep and MCF10A cells were cultured in DMEM/F-12 supplemented with 5% horse serum (Gibco), 20ng/ml epidermal growth factor, 0.5 μg/ml hydrocortisone, 100 ng/ml cholera toxin, 10 μg/ml insulin, and 1% pen/strep. To inhibit specific kinesins, cells were treated with 5 pmol siRNA with Lipofectamine RNAiMAX Transfection Reagent (Invitrogen) in Opti-MEM Reduced-Serum Media (Gibco). Specific siRNAs include pools of Silencer and Silencer Select KIF18A (Invitrogen), KIF18B (Dharmacon), KIF4A (Invitrogen), KID/KIF22 (Invitrogen), MCAK/KIFf2C (Dharmacon), MAD2 (Invitrogen), CLASP1 (Dharmacon), HSET/KIFC1 (Dharmacon), and pools of scrambled-sequence negative control siRNAs (Invitrogen). For double knockdowns involving the inhibition of two proteins, Lipofectamine RNAiMAX was used at a lowered concentration (0.7X the concentration used for single knockdowns) to mitigate toxicity.

### Generation of Inducible HeLa Kyoto Cell Line

HeLa Kyoto cells that inducibly express GFP or GFP-KIF18A were generated via recombination mediated cassette exchange, as previously described^49^. Briefly, a wild-type KIF18A siRNA and puromycin resistant plasmid was developed containing LoxP sites for recombination mediated cassette exchange. The LoxP containing plasmid was then transfected into HeLa Kyoto acceptor cells (a kind gift from Ryoma Ohi’s lab)^50^, and cells which had undergone recombination were selected via puromycin. The open reading frame for a KIF18A wild-type siRNA resistant construct^51^ and pEM791 vector^49^ were amplified with primers designed for Gibson Assembly (New England BioLabs). After confirming the correct sequence of the Gibson assembled plasmid, recombination was achieved by transfecting the acceptor cells with the KIF18A plasmid and recombinase using an LTX transfection (Thermo Fisher Scientific). Recombination was initially selected for with 1 μg/mL puromycin for 48 hours followed by a stricter selection with 2 μg/mL puromycin for 48 hours prior to switching back to 1 μg/mL puromycin. The resulting wild-type KIF18A inducible cell line was maintained in MEM Alpha (Life Technologies) with 10% FBS (Life Technologies) and 1 μg/mL puromycin at 37°C, 5% CO_2_.

### Drug Treatments

For experiments involving siRNA knockdown followed by drug treatment, the indicated concentrations of paclitaxel (Selleck Chemicals), nocodazole (Selleck Chemicals), and/or monastrol (Selleck Chemicals) were added to cells 24 hours after siRNA treatment. Three hours after drug addition, cells were either fixed and stained for immunofluorescence imaging or imaged live in a glass-bottom 24-well dish. To compare the effects of paclitaxel treatment to the effects of KIF18A KD in MDA-MB-231 and MCF10A cell lines, 10 nM of paclitaxel was added to cells 24 hours before fixing and staining for immunofluorescence imaging.

### Proliferation and Cytotoxicity Assays

Cells were imaged in either a 96- or 24-well dish every two or four hours for up to five days using the Cytation 5 Cell Imaging Multi-Mode Reader (Biotek) driven by Gen5 software (Biotek). A 4X Plan Fluorite 0.13 NA objective (Olympus) was used to capture images. Between imaging reads, cells were incubated at 37°C with 5% CO_2_ using the Biospa 8 Automated Incubator (Biotek). Gen5 software (Biotek) was used to process images and to measure cell confluence and the number of cells/mm^2^ using high-contrast brightfield images. Parameters including cell size and light-intensity thresholds were specified for each cell line. To determine rates of cell proliferation, the fold change in cells/mm^2^ between the first and last reads of each well were calculated and normalized to the control for each experiment. One-way ANOVA with post-hoc Tukey’s test was used to compare proliferation fold-change values across cell lines to determine statistical significance. For cytotoxicity assays, CellTox™ Green Dye (Promega) was added to cell media prior to imaging, and the number of cells/mm^2^ was recorded for both GFP and brightfield channels. After four days of imaging, the area under the proliferation curve for the CellTox-stained cells was divided by the area under the proliferation curve for the total number of cells, and this value was normalized to the control for each cell line as the metric for relative cell death. An unpaired t-test was used to determine significance between control and KIF18A KD for each cell line by comparing the normalized proliferation fold-change values.

### Automated Cell Count Validation

Cells were seeded in a series of increasing densities in either a 96- or 24-well dish and allowed to adhere for 24 hours. Cells were then incubated with Hoechst stain (Invitrogen), a cell-permeable nuclear dye, for 30 minutes before being imaged using the Cytation 5 system as described previously. For each field, one high-contrast brightfield image and one fluorescence image were acquired, and Gen5 software was used to process images and analyze the number of cells/mm^2^ using the parameters defined in the proliferation assays. The correlation between cell densities measured in the brightfield images and the fluorescence images was graphed as a scatterplot (Extended Data Fig 2).

### Immunofluorescence

Cells were grown on glass coverslips and fixed using either −20°C methanol or 1% paraformaldehyde in −20°C methanol. Cells were blocked with 20% goat serum in antibody diluting buffer (Abdil-TBS, 1% BSA, 0.1% Triton X-100, and 0.1% sodium azide) and incubated with the following primary antibodies: mouse anti-α-tubulin (DM1α) 1:500 (Millipore Sigma) for one hour at room temperature (RT), human anti-centromere antibody (ACA) 1:250 (Antibodies Incorporated) overnight at 4°C, rabbit anti-γ-tubulin 1:500 (Abcam) for one hour at RT, mouse anti-γ-tubulin 1:500 for one hour at RT (Abcam), rabbit anti-KIF18A 1:100 (Bethyl Laboratories) at 4°C overnight, mouse anti-centrin-1 1:500 (Santa Cruz Biotechnology) for one hour at RT, rabbit anti-mCherry 1:500 (Abcam) for one hour at RT, and rabbit anti-KIF18B 1:2000^52^ for one hour at RT. Secondary antibodies conjugated to Alexa Fluor 488, 594, and 647 (Molecular Probes) were used at concentrations of 1:15000 for one hour at RT. Coverslips were mounted onto glass slides using Prolong Gold anti-fade mounting medium with DAPI (Molecular Probes).

### Microscopy

Fixed and live cell images were acquired using a Ti-E or Ti-2E inverted microscope (Nikon Instruments) driven by NIS Elements software (Nikon Instruments). Images were captured using a Clara cooled charge-coupled device (CCD) camera (Andor) or Prime Bsi sCMOS camera (Teledyne Photometrics) with a Spectra-X light engine (Lumencore). For live-cell imaging, cells in CO_2_-independent media (Gibco) were imaged using Nikon objectives Plan Apo 20X 0.75 NA or 40X 0.95 NA and an environmental chamber at 37°C. Fixed cell images were taken using Plan Apo 40X 0.95 NA, Plan Apo λ 60× 1.42 NA, and APO 100× 1.49 NA (Nikon).

### Western Blot

Cells were lysed in PHEM lysis buffer (60 mM Pipes, 10 mM EGTA, 4mM MgCl_2_, and 25 mM Hepes) with 1% Triton X-100 and protease inhibitors, incubated on ice for 10 minutes, and centrifuged at maximum speed for 5 minutes. Laemmli buffer with β-mercaptoethanol was added to the supernatant prior to boiling for 10 minutes at 95°C. Lysates were run on 4-15% gradient gels (BioRad), transferred (75 minutes at 100V) to PVDF membrane (BioRad), and blocked for one hour in 1:1 Odyssey Blocking Buffer (Li-Cor) and TBS with 0.1% Tween-20. Membranes were incubated with primary antibodies overnight at 4°C. Primary antibodies included 1:1000 mouse anti-GAPDH (Invitrogen), 1:500 rabbit anti-KIF18A (Bethyl Laboratories), 1:1000 rabbit anti-Kif4A (Bethyl Laboratories), 1:1000 rabbit anti-KIF22 (Millipore Sigma), 1:1000 rabbit anti-MCAK (Abcam), 1:1000 rabbit anti-MAD2 (Bethyl Laboratories), and 1:1000 rabbit anti-Cleaved Caspase-3 (Cell Signaling Technology). Secondary antibodies included goat anti-Rabbit IgG DyLight 800 conjugate and goat anti-mouse IgG DyLight 680 (Invitrogen), which were each diluted to 1:15000 in 1:1 Odyssey blocking buffer/TBS and added to the membrane for one hour at room temperature. Membranes were imaged using an Odyssey CLx (Li-Cor).

### Live Imaging

Cells were plated in a glass-bottom 24-well dish and treated with the indicated siRNA approximately 24 hours before imaging. Six hours before imaging, the cell culture media was replaced with CO_2_-independent media containing 100μM SiR-tubulin (Cytoskeleton). For conditions involving UMK57 (a kind gift from Duane Compton) or DMSO, the specified drug was added to the CO_2_-independent media with siR-tubulin. To image centrosomes, MDA-MB-231 cells were transfected with RFP-pericentrin via a 4D nucleofector system (Lonza) prior to seeding in glass bottom dishes and siRNAs were added 24 hours later. Cells were imaged every 2 minutes for 16-20 hours using a 40X 0.75 NA objective (Nikon).

### KIF2C/MCAK Overexpression

MDA-MB-231 cells were transfected with either a construct containing mCherry fused to the N-terminus of full-length KIF2C/MCAK (mCh-MCAK-FL) or a construct containing the centromere binding domain of CENP-B fused to the C-terminus of mCherry and the N-terminus of MCAK (mCh-CPB-MCAK), both kind gifts from Linda Wordeman^41^. mCh-CPB-MCAK is similar to a previously described construct designed to increase MCAK activity at centromeres (RCPBM)^41^ but contains mCherry instead of mRFP and an additional eight amino acids linking CPB and MCAK to improve expression. Transfections were performed using a 4D nucleofector system (Lonza) and plated on to 12-mm glass coverslips. KIF18A or control siRNA was added to cells 24 hours post-transfection, and cells were fixed and stained for immunofluorescence after an additional 24 hours.

### Mitotic Timing and Mitotic Index Analyses

To measure the length of mitosis, live cells were imaged every two minutes for 16-20 hours using differential interference contrast (DIC) microscopy. The time between nuclear envelope breakdown (NEB) and anaphase onset (AO) was used to indicate the time a cell spent in mitosis. Mitotic index was measured using fixed-cell images by counting the number of mitotic cells divided by the total number of cells. All mitotic index fields were taken with a 40x objective. An unpaired t-test was used to determine statistical significance between control and KIF18A KD conditions for each cell line using the mean percentages of mitotic cells from each experimental replicate. Contingency tables were created to compare the total number of cells that either divided or failed to divide; statistical significance was then determined using Chi-square tests to compare between the conditions within each cell line. An unpaired t-test was used to determine the statistical significance between control and KIF18A KD conditions in the mitotic timing analysis by comparing the averages of all the timing values (minutes from NEB to AO) between the two conditions within each cell line. For HeLa Kyoto mitotic timing rescue experiments, a one-way ANOVA was used to compare averages of the timing values between the three conditions.

### Mitotic Spindle Morphology Analyses

To analyze mitotic spindle morphology, cells were fixed and stained for γ-tubulin, α-tubulin, and centrin-1. Enough optical slices spaced 200 nm apart were captured to visualize the entire 3-D structure of the spindle. Spindles with three or more visible γ-tubulin-containing microtubule-organizing centers were classified as multipolar. Contingency tables were created to compare the total number of bipolar and multipolar cells between conditions, and statistical significance was determined using a Chi-square test. To characterize the structure of microtubule organizing centers across different conditions, pericentriolar material (PCM) and centrioles were stained and imaged in fixed cells. Cells were considered to possess fragmented pericentriolar material if they had supernumerary poles observed via γ-tubulin staining but lacked centrioles (centrin-1 puncta) at one or more of the poles. An unpaired t-test was used to determine statistical significance between control and KIF18A KD conditions for PCM fragmentation measurements. Intercentriolar distance, or the distance in microns between two centrioles in a pair, was measured from the center of one centriole to the center of the adjacent centriole. One-way ANOVA with post-hoc Tukey’s test was used to compare intercentriolar distance measurements between conditions.

### Analysis of Live Spindle Pole Dynamics

Movies of MDA-MB-231 cells stained with SiR-tubulin were analyzed to assess the timing of multipolar spindle formation. If a cell entered mitosis with three or more visible microtubule organizing centers, it was considered to have begun multipolar. If a cell entered mitosis with two microtubule organizing centers but ended up with three or more by the end of the movie, it was considered to have split poles. Contingency tables were created to compare the total number of cells that split poles and the total number that remained bipolar between conditions, and statistical significance was determined using a Chi-square test. The same analysis was conducted to compare differences across conditions in the proportion of cells that began multipolar.

### Knockdown Quantification Analysis

The efficiency of siRNA-mediated kinesin knockdowns was measured via either quantitative western blot or immunofluorescence. ImageJ was used for all quantification. KIF18A knockdown efficiency in CRC cell lines was measured by comparing background-subtracted KIF18A fluorescence intensity in cells treated with control or KIF18A siRNA. In TNBC cell lines, KIF18B knockdown efficiency was measured by comparing background-subtracted KIF18B fluorescence intensity in cells treated with control or KIF18B siRNA. All other knockdown quantifications were determined by Western blot analysis. For MCF10A and MDA-MB-231 cell lines, the KIF18A knockdown efficiency was further analyzed at the RNA level by qRT-PCR.

### qRT-PCR

Total RNA extraction was carried out using RNeasy Mini Kit (Qiagen). Extracted RNA was screened by the Vermont Integrative Genomics Resource (VIGR) DNA Facility for purity and integrity using a 2100 Bioanalyzer (Agilent Technologies), and human GAPDH and human KIF18A Taqman probes and primers (Thermo Fisher Scientific) were used for reverse transcription and qRT-PCR. KIF18A RNA expression levels were normalized to GAPDH RNA levels in each cell line.

## Acknowledgements

The authors wish to thank Dr. Carol Vallett, Dr. Marion Thurnauer, Dr. Marie Wood, and Dr. Christopher Anker for insightful discussions and suggestions. We also thank Dr. Uri Ben-David and Dr. Neil Ganem for sharing unpublished results and Linda Wordeman and Duane Compton for reagents. This work was supported by Susan G Komen grant CCR16377648, NIH grant GM121491, American Cancer Society Institutional Research Grant 14-196-01, and a Lake Champlain Cancer Research Organization pilot grant (to JS) and by Department of Defense PRCRP Horizon Award W81XWH-17-1-0371 (to HM). The authors declare no competing financial interests.

